## Supplementary figures and images for "Late-life dietary folate restriction reduces biosynthetic processes without compromising healthspan in mice"

### Figure 7S1 - source sata

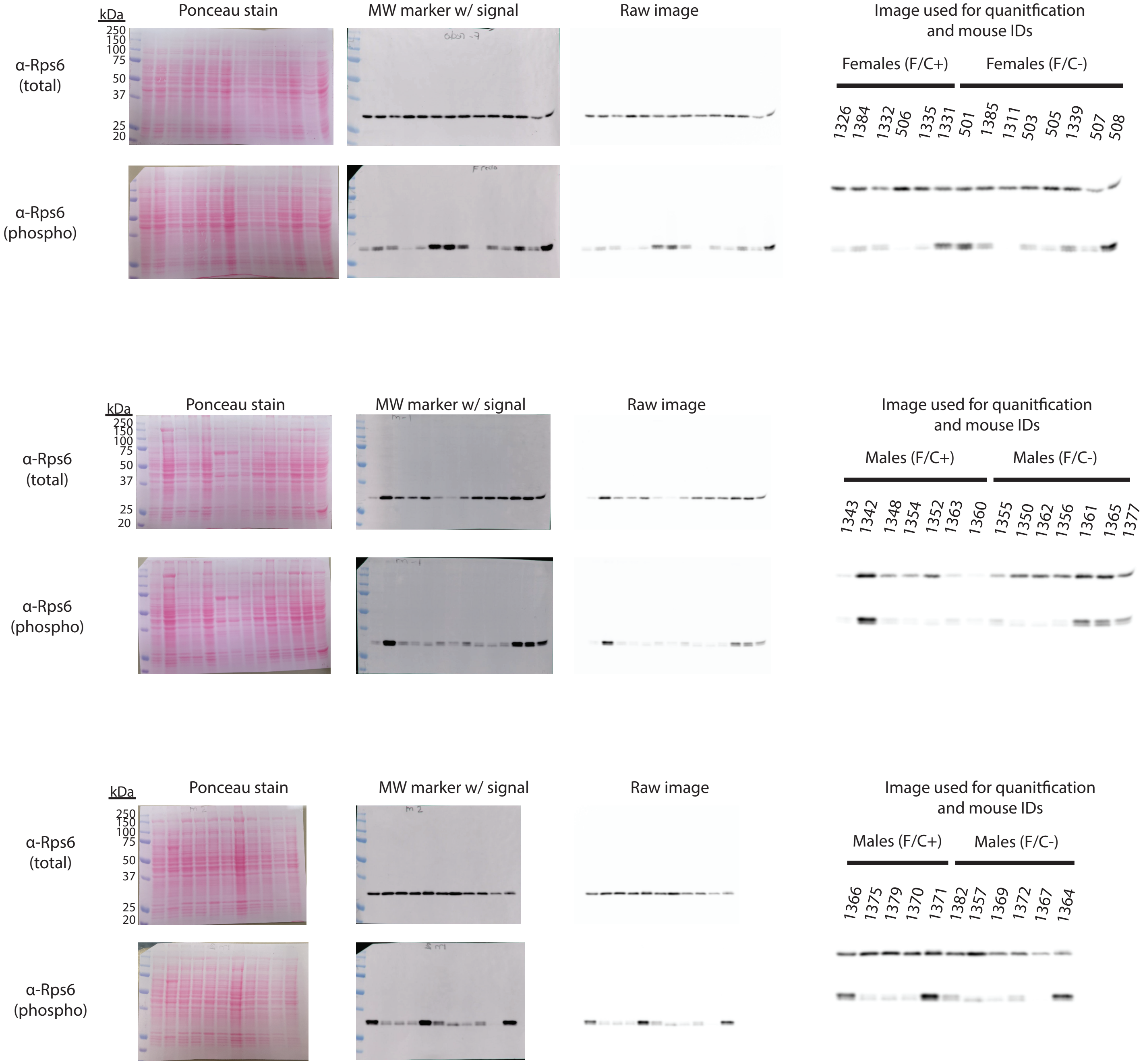
